# Adaptive Efficient Coding: A Variational Auto-encoder Approach

**DOI:** 10.1101/2020.05.29.124453

**Authors:** Guy Aridor, Francesco Grechi, Michael Woodford

## Abstract

We study a model of neural coding with the structure of a variational auto-encoder. The model posits that the encoding of individual stimulus values is optimally adjusted for a finite training sample of stimuli retained in memory. We demonstrate that this model can rationalize existing experimental evidence on both perceptual discrimination thresholds and neural tuning curve widths in multiple sensory domains. Finally, since our model implies that encoding is optimized for a sample from the environment, it also provides predictions about the adaptation of neural coding as the environmental frequency distribution changes.

## 1 Introduction

An influential literature has proposed that variation in the degree of precision with which sensory magnitudes are encoded over the range of possible stimulus values can be explained by a principle of *efficient coding* [3,4,25], according to which a finite range of possible internal representations is used in a way that is well-adapted to the frequency distribution of the stimuli that the organism encounters in its environment. A variety of assumptions have been proposed as to the precise formulation of the relevant constraint on feasible encoding schemes, and the performance measure that an efficient coding scheme should maximize; under one popular proposal (“infomax” theories), the efficiency criterion should be maximal mutual information between the internal representation and the objective stimulus magnitude [11, 12, 16, 17, 28].

A relatively neglected topic in this literature has been the question of how an efficient internal representation scheme is supposed to be learned from experience of instances of individual stimuli. This question is relevant both for understanding how cortical maps self-organize during development, and for understanding the speed and reliability with which an efficient encoding scheme for a new frequency distribution should be expected to arise when the statistics of the environment change. In this paper we propose a statistical learning approach to neural coding that draws on recent work in unsupervised representation learning.

We posit that the way in which internal representations of sensory stimuli are formed and used in the nervous system has the structure of a *variational auto-encoder* (VAE) [14, 15]. Such a system includes both an encoding circuit, that produces a low-dimensional internal representation for any presented stimulus, and a decoding circuit that can produce a reconstructed value for the original stimulus on the basis of this coarse categorization. The decoder learns a generative model of the stimulus distribution in the environment, in which the category to which a stimulus is assigned plays the role of a latent explanatory variable; the encoder is assumed to be optimized to label stimuli in a way that will make the labeled data useful for training the decoder. This architecture is useful, not only because it provides an arguably realistic model of what encoding schemes are adapted to do well, but because it provides a model of how unsupervised learning of a set of coarse categories appropriate to a given environment can occur.

We show that this model of neural coding produces predictions regarding the widths of neural tuning curves and discrimination thresholds that are consistent with evidence from multiple sensory domains, so that it is competitive with other proposed models of efficient coding (e.g.,[12]) in this respect. At the same time, our model provides an account of how an efficient coding scheme can be learned, and naturally allows the coding scheme to rapidly adapt to a new statistical environment, as we illustrate through a numerical example.

## 2 A Model of Neural Coding

### 2.1 General Setup

We suppose that each stimulus is described by a single real number *x* drawn independently from a continuous frequency distribution *π*(*x*). Each stimulus is to be encoded as belonging to one of *J* latent categories, 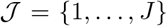; this bound on the number of possible categories is taken as a constraint. We consider neural coding systems with the structure of a VAE. In particular, we suppose that the nervous system learns an encoding rule that stochastically classifies a continuous stimulus as belonging to one of the discrete set of latent categories, with probabilities *p*(*j*|*x*), and a decoding rule that stochastically decodes the latent category back to a continuous stimulus magnitude, with probabilities 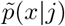. Thus the perceived stimulus 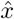 is the stochastic output of the original input being encoded according to *p*(*j*|*x*), and then decoded back to the stimulus space according to 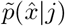. The collection of distributions 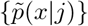, together with learned frequencies of occurrence {*q*(*j*)} of the latent categories, form a generative model for the distribution of stimulus magnitudes in the environment; the distribution 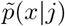 can be thought of as a “posterior” distribution for the stimulus magnitude when a given stimulus is encoded using category *j*, in a model of approximate inference.

The encoding rule, or recognition model, must be chosen from a parametric family of possible rules, *p_ϕ_*(*j*|*x*), where *ϕ* is a finite-dimensional vector of parameters. The encoding rule, combined with the environmental distribution *π*(*x*), implies a joint distribution for true stimulus magnitudes and their labels given by

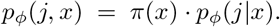

The decoding rule is likewise chosen from the family of parametric models, *p_θ_*(*x*|*j*), where *θ* is a finite-dimensional vector of parameters. The implied generative model for the joint distribution of stimulus magnitudes and labels is then given by

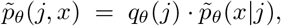

where we include the frequencies {*q*(*j*)} among the elements of *θ*. The problem for the recognition model is one of inferring the latent category *j* that has given rise to stimulus *x*, according to the generative model.

We can interpret this as a model of neural coding in a region of sensory cortex (say, the visual cortex), under the theory that the cortex maintains an internal generative model of how images are generated by underlying visual features, and that the role of early processing (for example, in V1) is then to invert this generative process and infer the extent to which a given image contains each of the possible features [18,21,23]. Following [9,19], we suppose that each of our categories *j* represents a possible feature, and corresponds to a particular population of cortical neurons, with the rate of firing in each of the populations indicating an inferred posterior distribution over the possible features in the image. Under this interpretation, the conditional probabilities *p*(*j*|*x*) in our model correspond to the relative firing rates of *J* populations of neurons. This allows us to derive quantitative predictions about the distribution of neural tuning curves, in addition to the model’s predictions about the discriminability of different stimuli.

We suppose that the neural coding system is organized to learn a good representation of the environment. But as we do not assume that it is optimized for a particular downstream task, it remains a question what objective function the coding system ought to optimize; there has been considerable debate in the representation learning literature about which objectives lead to the most useful representations [5, 26]. A natural approach would be to follow [14, 15] and suppose that *ϕ* and *θ* are jointly optimized so as to minimize the Kullback-Leibler divergence of the joint distribution implied by the encoder relative to that implied by the decoder, 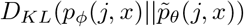. However, as noted by [7, 8], this would ensure a reasonable approximation to the environmental distribution *π*(*x*), but would not necessarily lead to a meaningful latent representation. Instead, we follow [2], who propose extending the objective function used in [14, 15] to explicitly incentivize the model to learn a more meaningful representation. Their “*β*-VAE” approach allows for more “disentangled” representations by providing an additional bonus for classification schemes in which the different categories are more informative about the underlying stimuli (the objective proposed in “infomax” theories).

Formally, we suppose that the parameters are optimized to solve the problem:

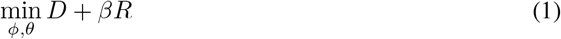

*D* is a measure of the average distortion and *R* is a measure of the complexity resulting from the coding scheme. *β* trades off the relative importance assigned to minimizing distortion as opposed to complexity where a lower *β* implies a higher relative importance assigned to having informative categories. *D* and *R* are defined formally as follows:

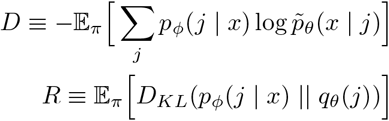

We suppose there is no restriction on the choice of {*q_θ_*(*j*)}. With unrestricted choice then it is clearly optimal, regardless of the value of *β*, to choose *ϕ* and *θ* such that *q_θ_*(*j*) = *p_ϕ_*(*j*). Under this condition, (1) reduces to:

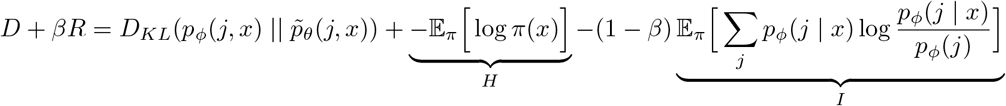

*H* represents the underlying entropy of the stimuli and is independent of *ϕ* or *θ*. *I* represents the Shannon mutual information between the category *j* and state *x* in the joint distribution produced by the recognition model. Thus, when *β* < 1, this objective assigns an additional bonus to classifications with higher mutual information between *j* and *x*, as desired. The smaller is *β*, the greater the emphasis placed on having more informative categories.

### 2.2 Training Data

We suppose that the parameters of the coding scheme are fit to a finite sample of observations drawn from the environment. Given a large set of previously observed stimuli, a sampling process selects a corpus of observations to be used in training the algorithm. We adopt a version of *reservoir sampling*, a standard algorithm from stream-processing [27], that provides a guarantee that, after any number of observations has been drawn, every previously observed stimulus has the same probability of being in the sample. Crucially, the standard version of reservoir sampling requires no knowledge of the total number of observations to be drawn and has a memoryless insertion and deletion policy, requiring only the current observation and the previous sample to be stored in memory at any given point.

We utilize a modification of traditional reservoir sampling, proposed in [1], that places greater weight on more recent observations over older observations. This temporal bias enables more rapid adaptation, as it allows for the possibility that the underlying environmental statistics are changing without requiring that the organism be explicitly aware of this shift. We utilize the version of reservoir sampling in [1] that results in an exponential bias in the sampling procedure but maintains the same desirable memoryless insertion and deletion policy. In particular, the algorithm has the property that the probability that the *r*-th observation being in the sample after the *t*-th observation is given by *f*(*r, t*) = *e*^-λ(*t−r*)^. In this case, 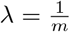 where *m* is the overall memory size. Thus, we introduce a single additional parameter specifying the memory size *m* which will implicitly define the degree of temporal bias.^1^

### 2.3 Parameterization and Learning Process

We illustrate our approach using a parameterization in which the generative model must be a finite mixture of Gaussians. In particular, we assume that for each 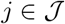:

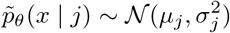

The parameters *θ* then consist of values {*q_j_, μ_j_, σ_j_*} for each 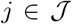, where the *q_j_* represent the mixture coefficients. We further assume that the parametric family of possible recognition rules is optimally adapted to this family of generative models, in the sense that for any generative model *θ*, there exists a recognition rule *ϕ* = *θ* that minimizes *D* + *βR* (over all possible recognition rules).

Thus our parametric family of recognition rules is given by

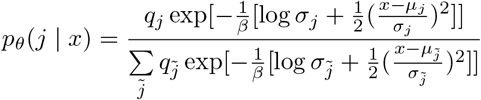

where the possible values of *ϕ* correspond to possible values of *θ*. Note that this kind of recognition rule can be implemented by a competition between *J* populations of neurons, in which the probability of a neuron in population *j* firing first (resulting in classification of the stimulus as of type *j*) in the case of stimulus *x* is proportional to the height of tuning curve *j* at point *x* in the stimulus space, and the tuning curves are Gaussian in shape. With this interpretation, our model makes predictions not only about the discriminability of different stimuli, but also about the distribution of preferred stimuli and tuning curve widths in a neural population code.

The problem of fitting the parameters of our model to a training data set reduces to the familiar problem of fitting the parameters *θ* of a Gaussian Mixture Model, with the small modification that we minimize *D* + *βR* rather than maximizing the likelihood. We use a training data set that is generated according to the procedure in subsection 2.2, and utilize an Expectation-Maximization (E/M) algorithm to fit the parameters of our model. For the implementation of the E/M algorithm we follow [6], but with modifications to the likelihood function that are required by the inclusion of *β* in *p_ϕ_*(*j*|*x*). Note that when *β* = 1, our procedures become identical to those described in [6].^2^

## 3 Stimulus Encoding in a Stationary Environment

We first consider a stationary environment where the underlying stimulus distribution is fixed, and consider the results from our model when 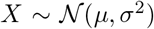 with *μ* =1 and *σ* = 1, though the qualitative patterns we identify hold for other distributions.^3^ Furthermore, we set *m* = 10,000 and show the results for this fixed memory size.

### 3.1 Properties of the Learned Model

In this section we describe the qualitative properties of our neural coding model as we vary *β* and *J*. We first fix *J* =10 and document the qualitative differences that result from varying *β*, and then proceed to analyze the case when *J* also varies. The qualitative patterns that emerge when varying *β* are robust to variation in *J*, and lead to significant qualitative differences in the resulting coding scheme.

We consider the grid of *β* ∈ {0.5,0.75,0.99}. The top row of Figure 1 shows the marginal distribution for *x* implied by the learned generative model (the VAE’s approximation of *π*). Note that, as expected, when *β* gets closer to 1 the resulting marginal closely approximates the true *π*. As we lower *β*, the resulting marginal is a worse approximation to *π*. But this is expected, as a lower *β* leads the objective function to place more weight on having meaningful latent representations and less weight on ensuring a close approximation to the true *π*.

**Figure 1:**
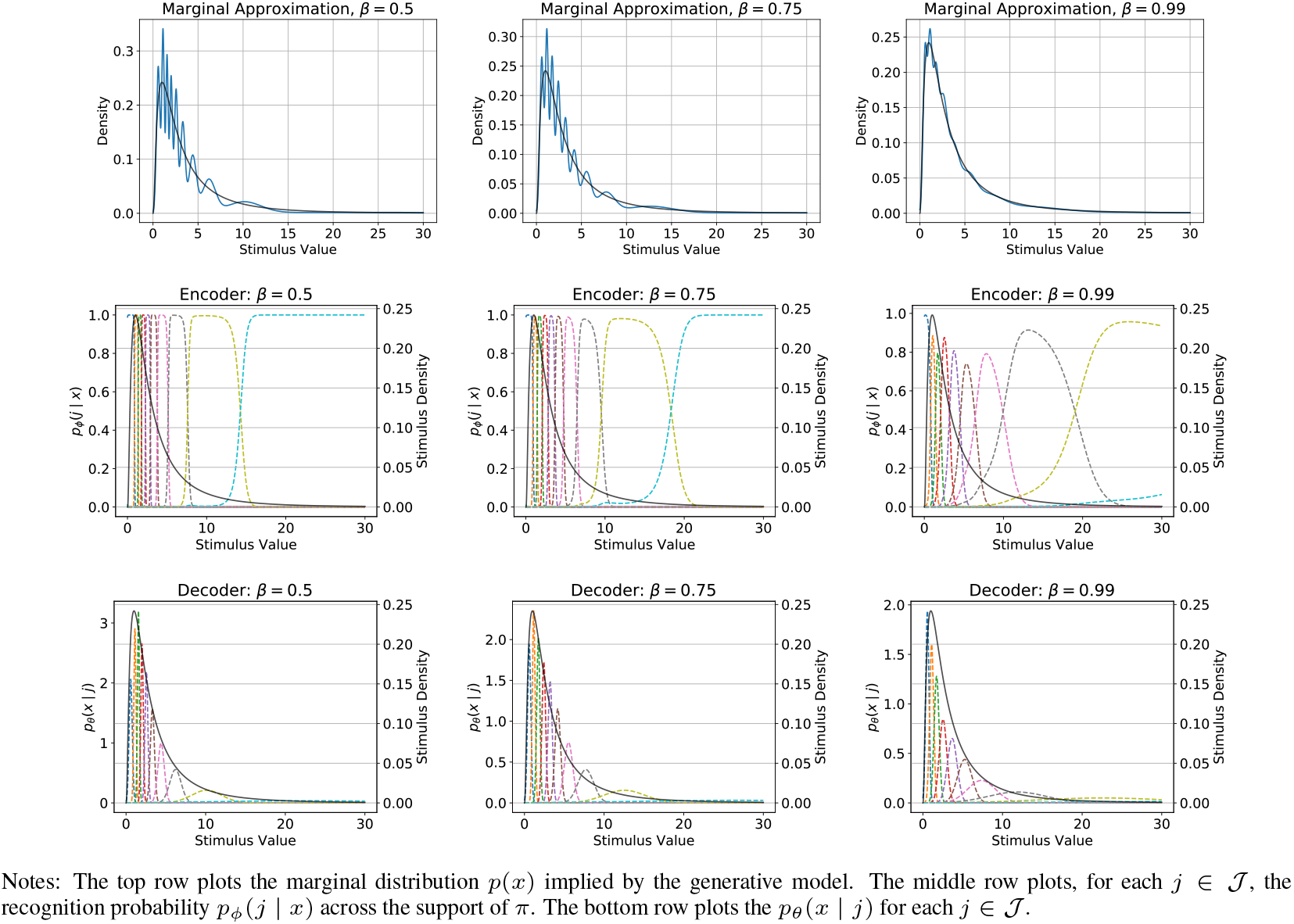
Marginal Approximation and Encoder / Decoder for *J* = 10, varying *β*

The middle and bottom rows of Figure 1 trace out *p_ϕ_*(*j* | *x*) and *p_θ_*(*x* | *j*), respectively, across all 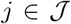 and *x* ∈ *supp*(*π*). Several qualitative patterns are apparent. The first is that, contrary to models of neural coding where tuning curves are shifted versions of the same function [20,24,30,31], in our model this is not generally the case. Instead, our model predicts that in regions of the stimulus space with high probability density, there is a higher density of neurons with narrower tuning curves whereas in portions of the stimuli space with lower probability mass there is a lower density of neurons with wider tuning curves.

There are also notable differences between the coding schemes obtained for different values of *β*. As *β* decreases, the encoding rule is increasingly deterministic, as is clear from Figure 1 and further apparent in Figure 2. Indeed, for *β* = 0.5 we find that the encoder is nearly deterministic except at stimulus values that are near category boundaries. However, for *β* = 0.99, there is considerable stochasticity in the optimal encoder.

**Figure 2:**
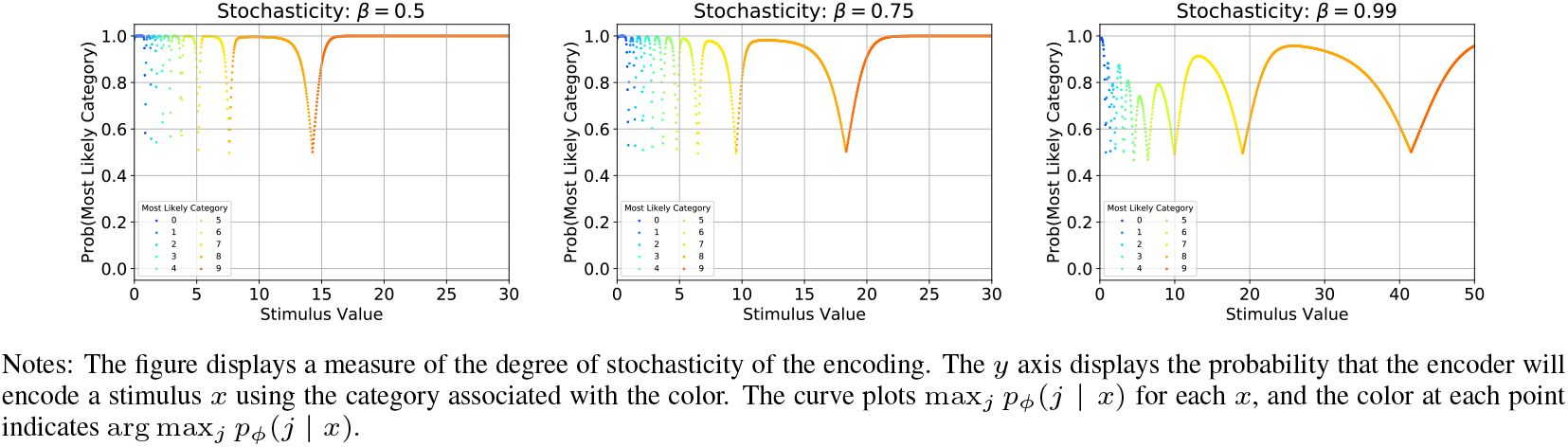
Degree of Stochasticity for *J* = 10, varying *β*

Finally, we investigate the role that *J* plays in the resulting coding scheme. Sufficiently low values of *J* lead to poor approximations of *π*, even in the case that *β* =1. Furthermore, numerical experiments confirm that increasing *J* weakly increases the value of the objective function. A natural question is how the coding scheme changes as we vary *J*. One possibility might be that as we increase *J*, the standard deviation of the various components does not change, but instead the components increasingly overlap. However, Figure 3 shows that as we increase *J* or decrease *β*, the mean *σ* and tuning curve width for the resulting models decrease. Thus, rather than having the same width components tiled across the stimulus space more densely, the components become narrower as *J* increases, so that their degree of overlap does not greatly increase.

**Figure 3:**
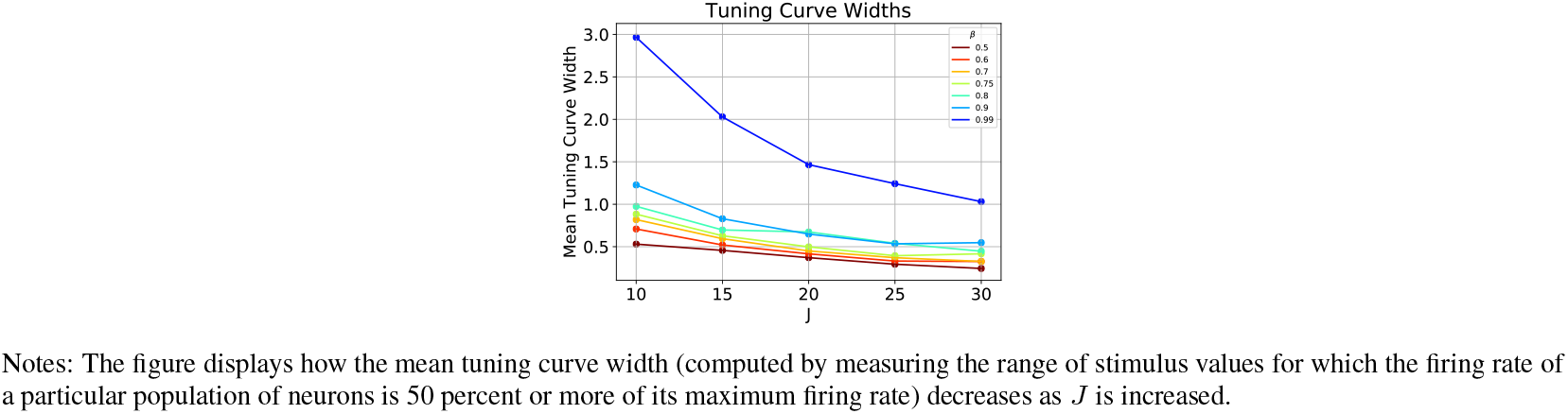
Tuning Curve Widths

### 3.2 Discriminability

In this section we apply our model of neural coding to generate predictions about stimulus discriminability. We characterize the qualitative differences in predictions as we vary *β* and *J*. In order to be qualitatively consistent with existing experimental evidence, our model should predict that the ability to discriminate between stimuli is inversely proportional to the frequency of occurrence associated with this stimuli in the environment [12].

In order to study discriminability, we can define a *just noticeable differences* (JND) for each stimulus value *x*:

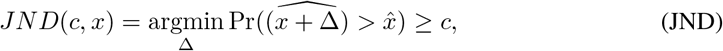

where 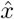 is the random category to which a stimulus *x* is assigned, and c > 0.5. Figure 4 displays the resulting JND values for *c* = 0.71,^4^ along with the density function *π* for comparison. It is apparent that regardless of the value of *β*, the qualitative predictions from the model are in line with experimental evidence, in that the predicted discrimination thresholds are lower for stimuli that occur more frequently in the environment.

**Figure 4:**
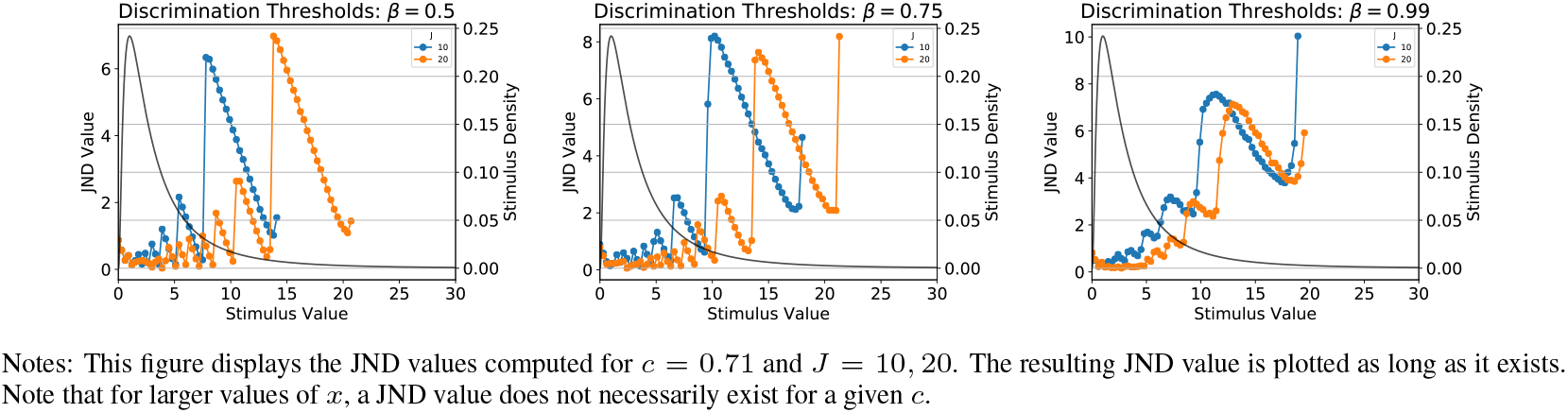
Discrimination Thresholds for *J* = 10, 20, varying *β*

We observe in Figure 4 that as *β* increases, the JND values become smoother. Indeed, for lower *β* values we find a sawtooth pattern in the JND values, owing to the small overlap between successive components in this case, as already noted. Differentiation between two different stimuli requires that they be encoded as belonging to a different category; thus when categories are relatively discrete, discrimination thresholds are highest at the lower boundary of a category and gradually decrease as the upper boundary is approached. When the stimuli begin to be mapped into the next category, there is a jump in the threshold, leading to the sawtooth pattern.^5^ Increasing *β* increases the JND curve’s smoothness because it increases the amount by which the tuning curves overlap. This also reduces the model’s ability to discriminate between nearby stimuli and thus results in larger JND values.

We also see that JND are decreasing in *J* for fixed *β*. Even for *β* = 0.5, we observe the same qualitative pattern for both *J* =10 and *J* = 20, but the JND values are lower when *J* = 20. The mechanism behind the sawtooth pattern is the same, except that, as shown in Figure 3, the components are more concentrated as *J* increases. Increasing *J* narrows the width of the categories, resulting in smaller JND values.

## 4 Calibrating the Model to Experimental Evidence

In this section we compare the predictions of our model to an empirically observed stimulus distribution, the modulation frequency distribution reported by [12]. This is estimated from a compilation of animal vocalizations, background sounds, and recordings made while walking around a suburban university campus. Furthermore, [12] compile empirical evidence from existing studies of neural tuning widths and discrimination thresholds of organisms in this environment.^6^

We calibrate *β* and *J* to illustrate that the model can rationalize the experimentally observed discrimination thresholds. We choose *β* and *J* to solve the following problem:

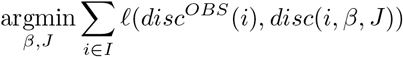

Here *I* denotes the set of experimental stimulus values, *disc^OBS^*(*i*) is the measured discrimination threshold for stimulus value *i*, and *disc*(*i, β, J*) is the predicted discrimination threshold for stimulus *i* given parameter values *β* and *J*. The loss function *ℓ* penalizes discrepancies between the two values, and we consider three possible specifications.^7^ Since it is computationally expensive to train the model for particular values of *β* and *J*, we search only over a discrete grid of possible values. We consider *β* in 0.01 increments from [0.5,0.95] and *J* ∈ {10,15, 20,25,30,35}. We pool the experimental results from the two separate studies in order to form the set I and the values *disc^OBS^*(*i*).

The resulting best fitting parameters for all three specifications of *ℓ* are *β* = 0.62 and *J* = 25. The observed and predicted discrimination thresholds as well as the underlying stimulus distribution are displayed in Figure 5. Figure 5 displays the results with the stimulus distribution presented in both logs and in levels, and shows that the calibrated model provides a reasonable fit to the measured discrimination thresholds in both experiments.

**Figure 5:**
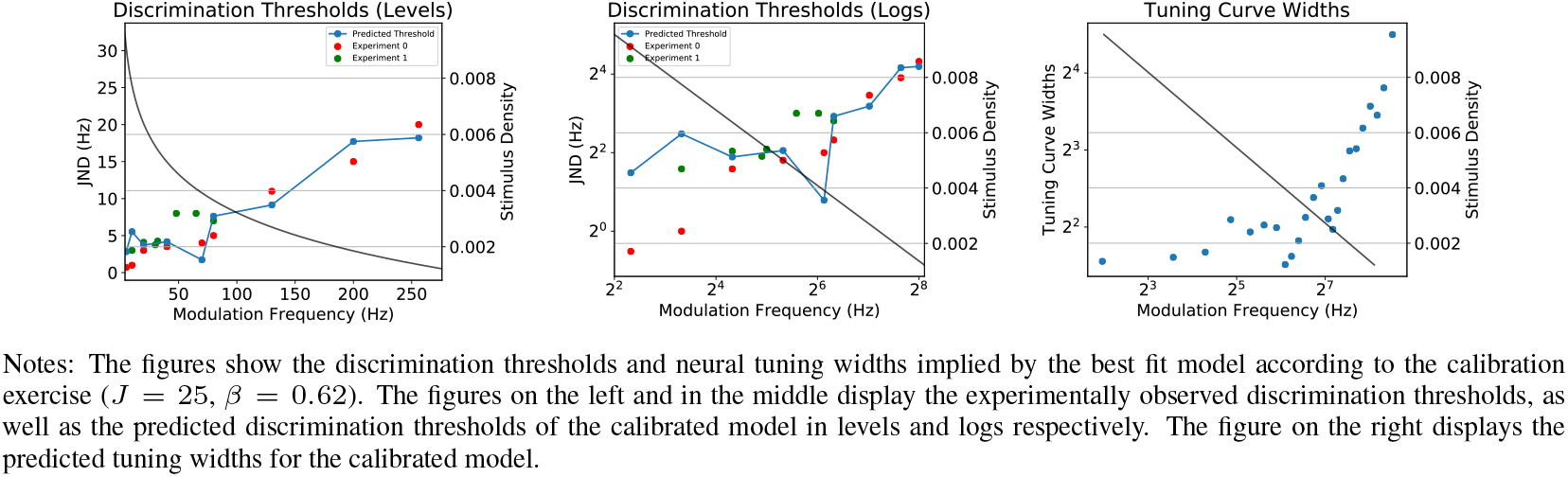
Discrimination Thresholds and Tuning Widths of Calibrated Model

We further investigate whether the predicted tuning widths of the calibrated model are consistent with physiological data. While it is difficult to directly compare the predicted tuning widths to the physiological data, we verify that our model produces predictions that are qualitatively consistent with the observed data. In particular, we expect that the tuning widths should be approximately inversely proportional to the probability density at the “preferred stimulus” of the particular tuning curve. We define the width of the *j*th tuning curve as the length of the interval of stimulus values for which the tuning curve amplitude (firing rate) is at least half the amplitude at the peak (see the supplementary material for details). The resulting predicted tuning widths are displayed in Figure 5 and match these patterns. Overall, the model’s predictions are in line with the experimental data on both discrimination thresholds and neural tuning widths.

## 5 Adaptation to a New Stimulus Frequency Distribution

In this section we demonstrate that the model also provides predictions about *adaptation* of the neural population code to a new environment. As an illustration, we consider a transition from *π*_1_ = log *X*_1_, 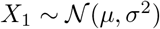, to 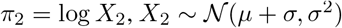, with *μ* =1 and *σ* = 1 as before.

Figure 6 displays the adaptation of *μ*, *σ*, and the marginal distribution for *x* implied by the learned generative model, where the first 25,000 samples are drawn from *π*_1_, the remaining 105,000 samples are drawn from *π*_2_. The parameters of the model are eventually fully adapted to *π*_2_ within this time period. The rate of convergence depends crucially on the memory size *m*, which is set to the same value as before (*m* = 10,000). In the supplementary material, we also present results for *m* = 1,000 and show that while the coding scheme adapts more rapidly with this lower memory size, it also induces additional jitter in the parameter values. Convergence occurs once the empirical distribution used to train the VAE fully transitions from a sample from *π*_1_ to a sample from *π*_2_, and this occurs faster with a lower *m*; but smaller *m* also leads to a less precise approximation of the distribution sampled from.^8^

**Figure 6:**
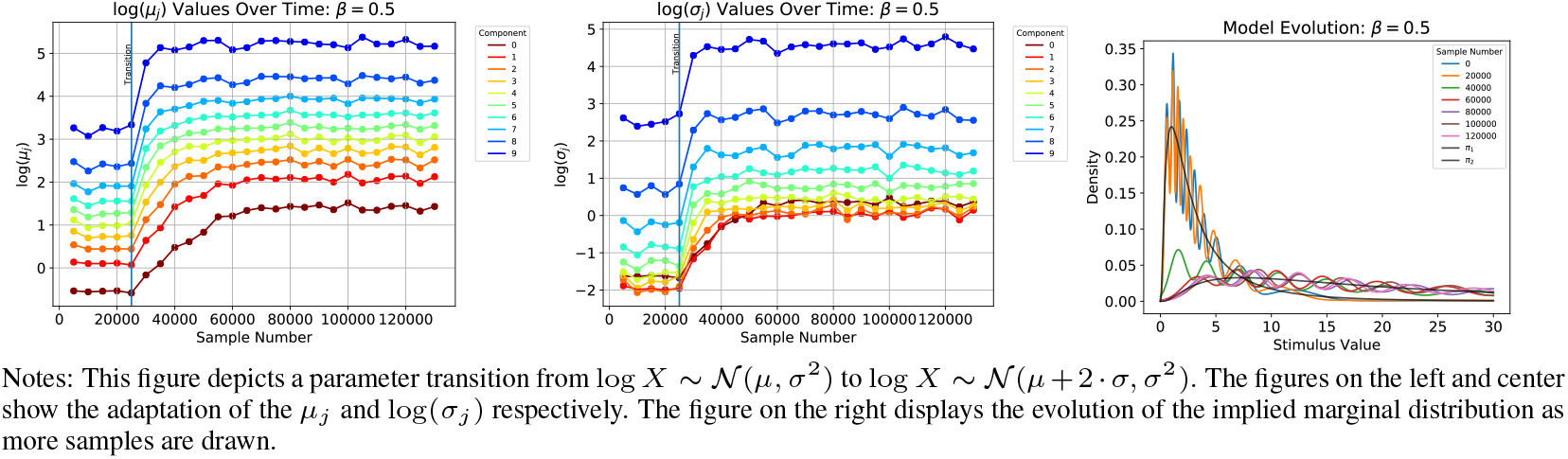
Adaptation of Parameters and Model

A key advantage of our model of adaptation is that at no point does the perceptual system need to be instructed that the environment has changed. Rather, the sampling method that determines the training data set is constructed to rapidly adapt to a new environment owing to the temporally biased sampling. In future work we hope to explore how the quantitative predictions of such a model of adaptation in a neural population code match empirical evidence.

## Appendix

## 1 Figures for Alternative Stimuli Distributions

In this section we demonstrate that the qualitative insights from Section 3 are robust to the choice of the stimulus distribution. As an alternative, we here we consider the case *π* ~ *Exp*(1.0) and reproduce the figures in Section 3 in the case of this alternative distribution.

**Figure 1:**
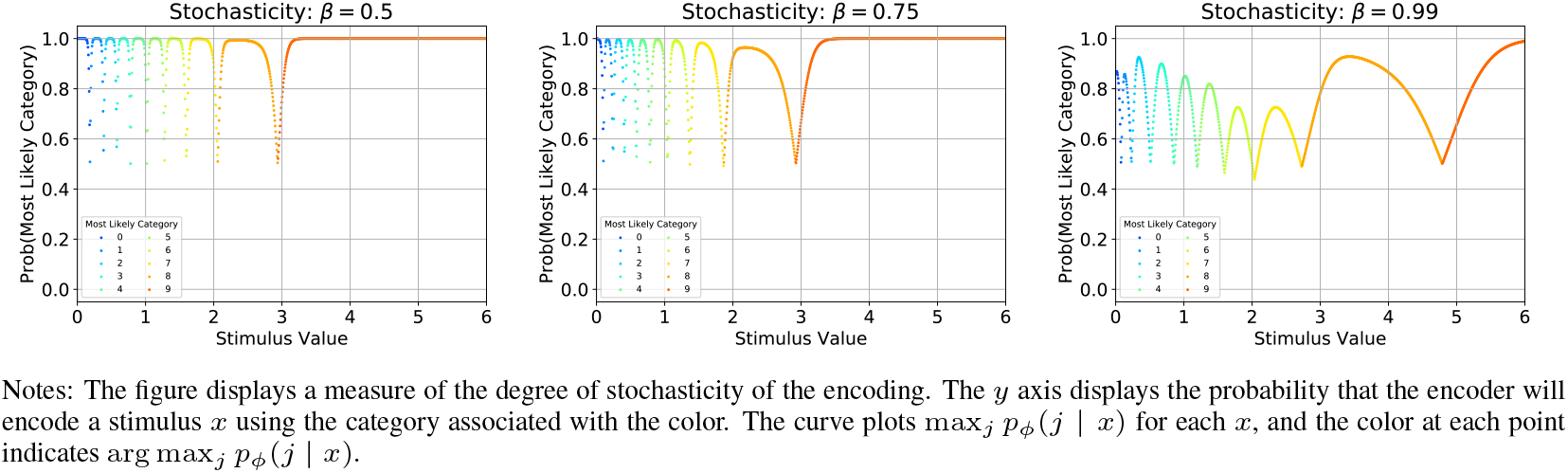
Degree of Stochasticity for *J* = 10, varying *β*

**Figure 2:**
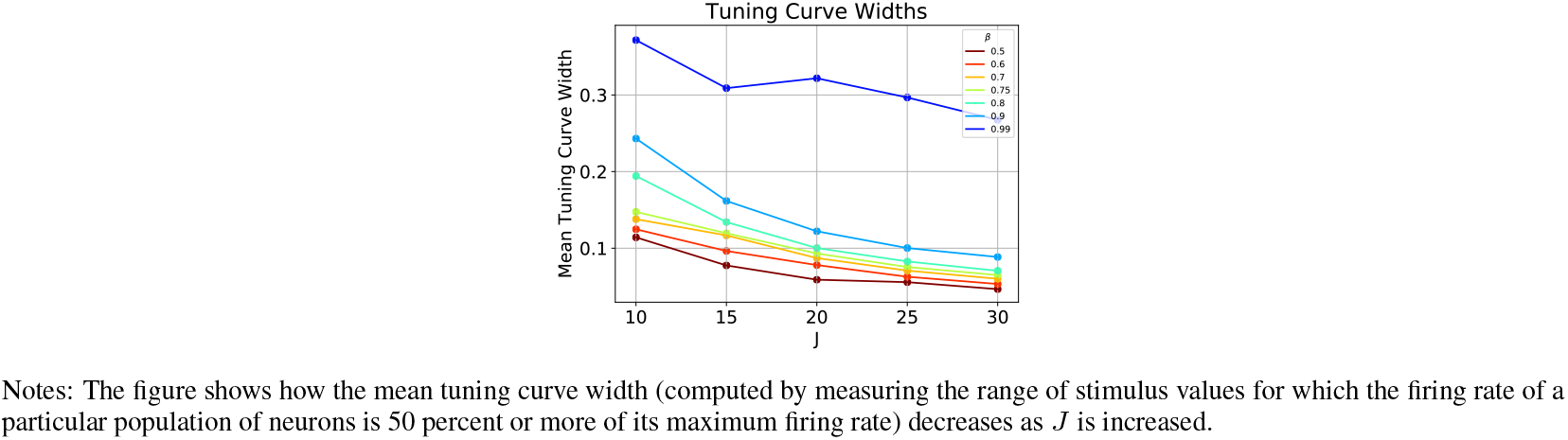
Tuning Curve Widths

**Figure 3:**
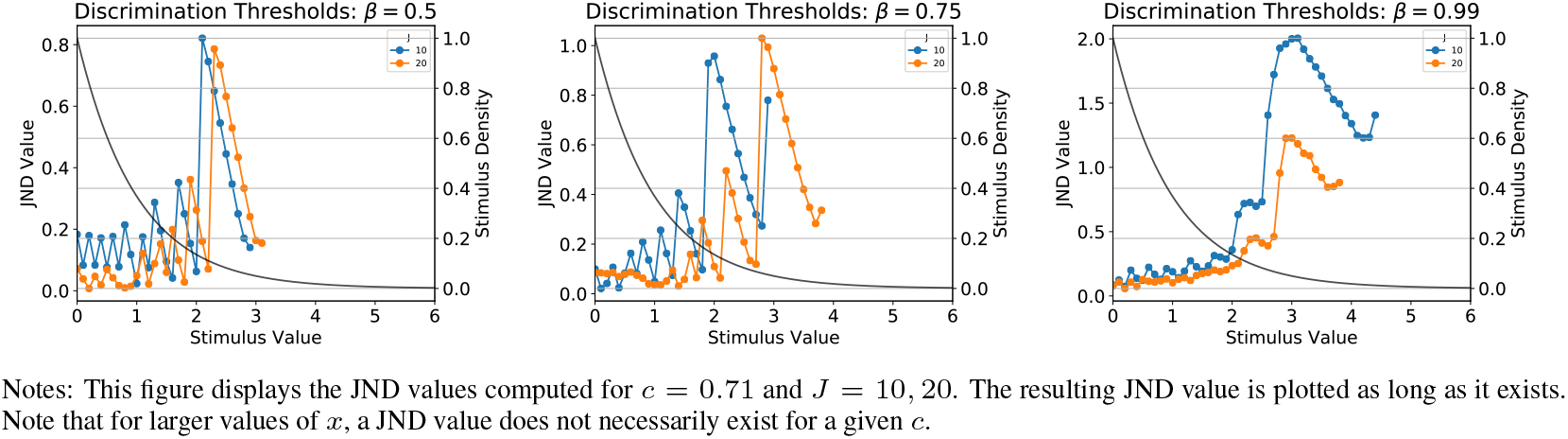
Discrimination Thresholds for *J* = 10, 20, varying *β*

**Figure 4:**
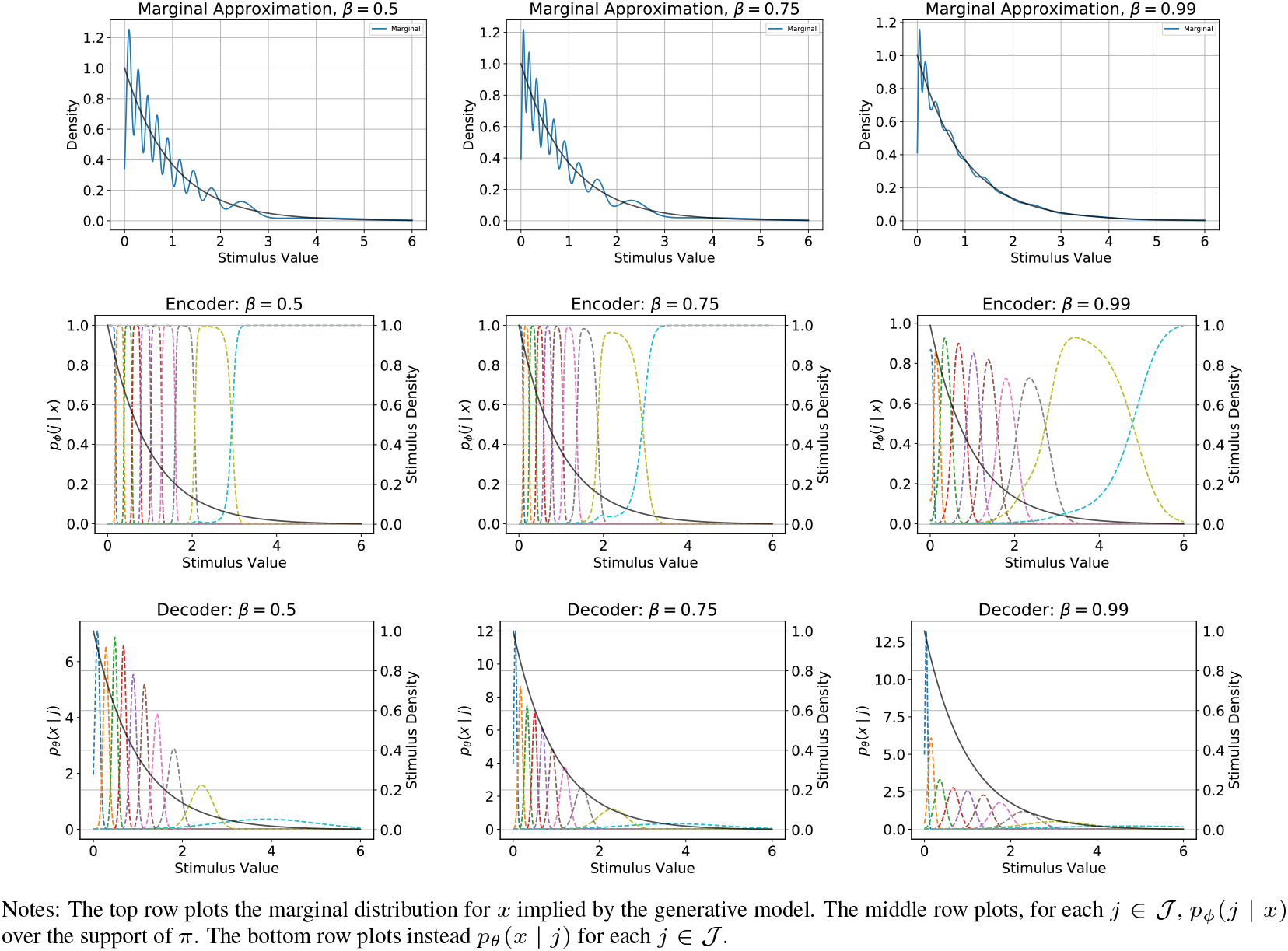
Implied Marginal Distribution and Encoder / Decoder for *J* = 10, varying *β*

## 2 Sample Adaptation

In this section we provide more details of the sampling procedure utilized to train the VAE in our model, and illustrate it via numerical examples. Algorithm 1 describes the details of the sampling algorithm from [1], a reservoir sampling algorithm with an exponential temporal bias. The algorithm guarantees that the probability that the *r*-th observed stimuli value is still present in the sample after the *t*-th observation is given by *f*(*r, t*) = *e*^−λ(*t−r*)^, where 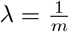 for a fixed memory size *m*. We focus on the case of adaptation following a transition from an original distribution *π*_1_ to a new distribution *π*_2_.

We first investigate the rate of decay of samples from the original *π*_1_ distribution. Figure 5 displays the fraction of remaining observations from *π*_1_ as more samples are drawn from *π*_2_. The figure confirms that there is an exponential decay of samples from *π*_1_, and that the rate of decay is faster for lower memory sizes. In the adaptation exercise in the main text, we use the value *m* = 10,000, and find that after 40,000 samples from *π*_2_ the samples from *π*_1_ have nearly vanished. We further demonstrate the resulting adaptation of the empirical distribution when 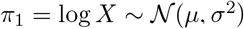 and 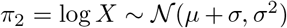 as well as 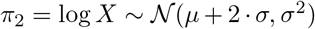 with *μ* = *σ* = 1.0. Figure 6 displays the resulting empirical distribution during the transition between *π*_1_ and *π*_2_.

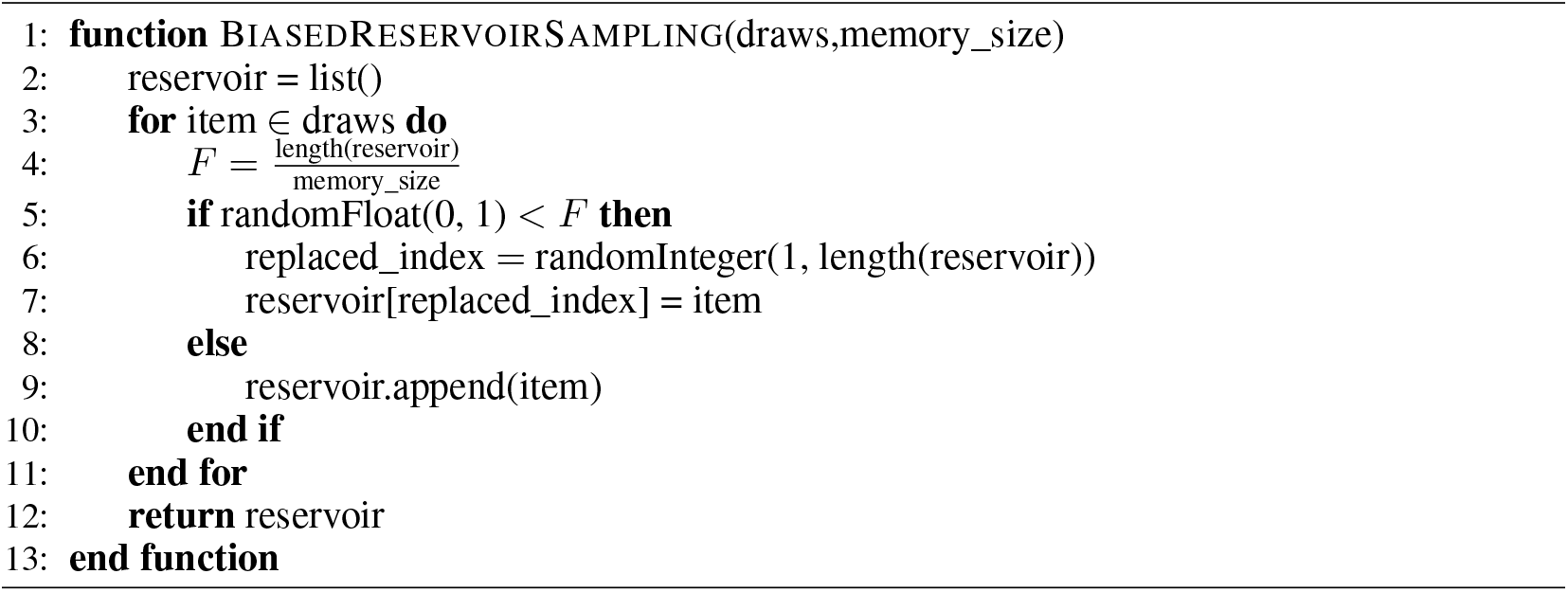

**Figure 5:**
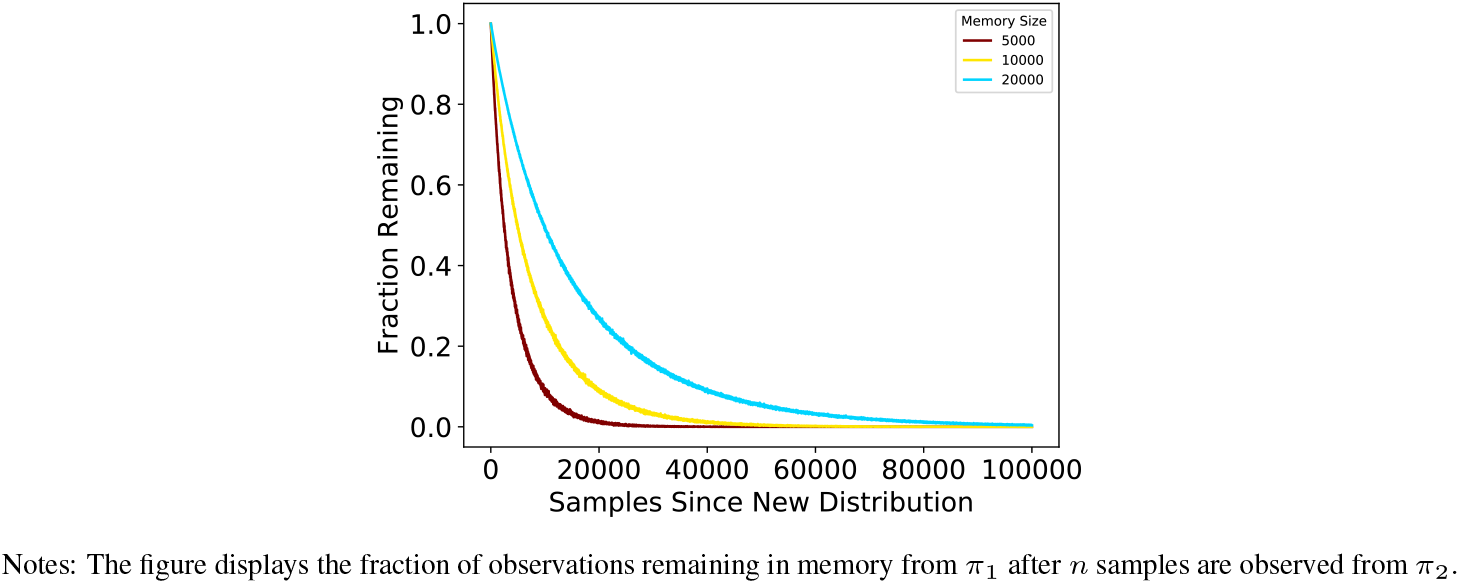
Rate of Decay of Observations from *π*_1_ in Memory

**Figure 6:**
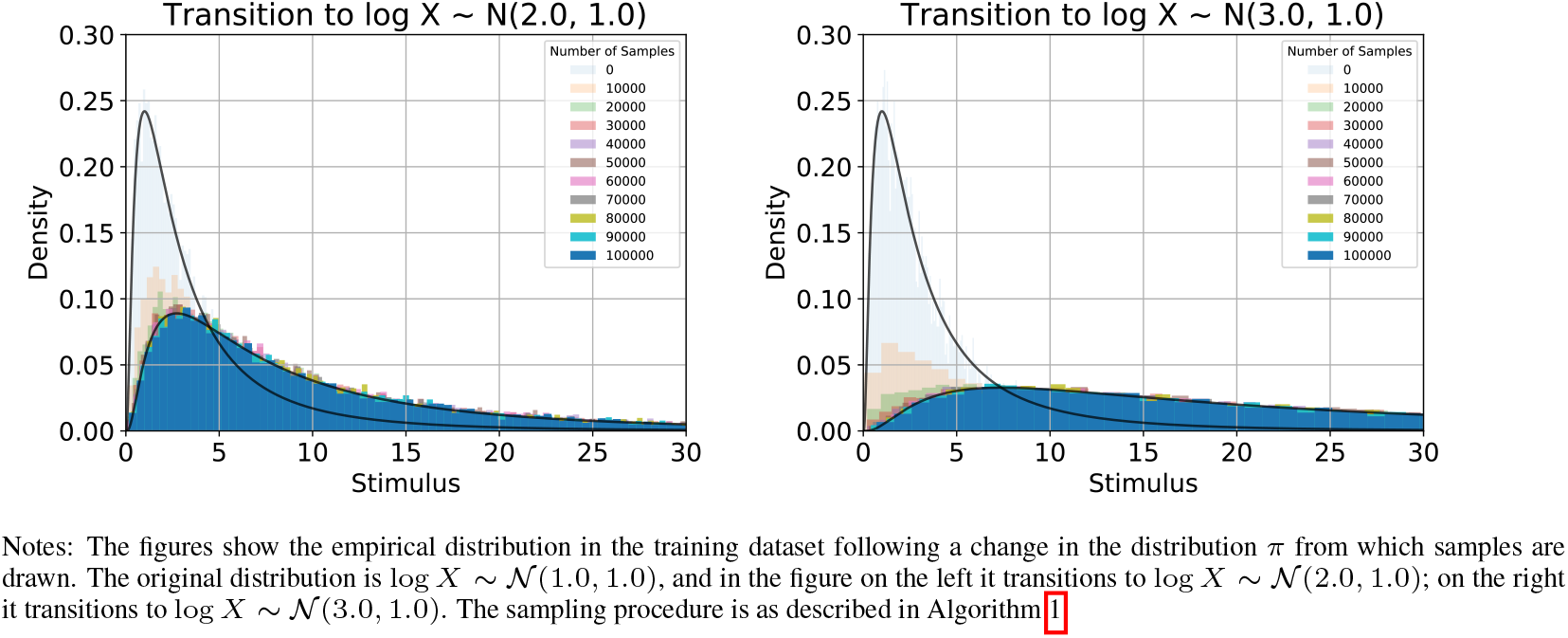
Empirical Distribution of Transition for *m* = 10,000

## 3 Neural Tuning Curve Computation

In this section we provide additional details on the definition of neural tuning curves in the context of our model and provide a closed form calculation for the resulting neural tuning curve widths that are described in the main text.

### 3.1 Parametric families of functions used

We consider a finitely parameterized family of possible generative models,

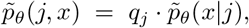

where the {*q_j_*} are also part of the vector *θ*. For any such generative model, the recognition model *ϕ* that will minimize *D* + *βR* (allowing for a completely flexible recognition model) will be one such that

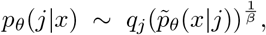

where we use the notation *ϕ* = *θ* since the parameters of this model are same as those of the generative model for which the recognition model has been optimized. Note that if *β* = 1, the recognition model assigns conditional probabilities *p_θ_*(*j*|*x*) that are equal to the conditional probability of a stimulus *x* having been produced as a draw from category *j* of the generative model, in the way that generative models are commonly used in Bayesian models of perception. However, when *β* ≠ 1, this is no longer exactly the case.

In the case that the family of generative models considered is the family of finite mixtures of Gaussians, then the parameters are *θ* = {*q_j_, μ_j_, σ_j_*}, and we have

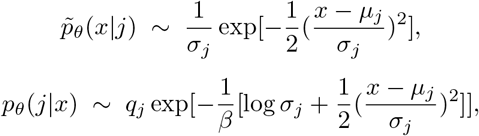

where in each case we have suppressed the common multiplicative factor required to make the conditional probabilities sum (or integrate) to 1.

### 3.2 A Neural Coding Interpretation

The recognition model *p_ϕ_*(*j*|*x*) can be implemented by competition between pools of neurons in the following way. Suppose that there are *J* pools of neurons, with *n_j_* neurons of each type *j*, and let *x* be a number on the real line indicating the physical magnitude of some stimulus feature. When a stimulus *x* is presented, each neuron of type *j* spikes at a Poisson rate proportional to *g_j_*(*x*), where the non-negative function *g_j_*(*x*) is the “tuning curve” for neurons of type *j*. We suppose that the stimulus is categorized as belonging to category *j* (i.e., is encoded by *j*) if the first spike is produced by a neuron of type *j*. Thus the recognition model implemented by the neural population is of the form

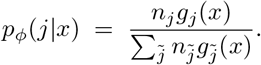

The parametric family of recognition models that we assume in our version of a *β*-VAE are of this form, where

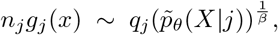

and *θ* indicates the generative model for which the recognition model has been optimized. In the case that the generative model *θ* is a mixture of Gaussians parameterized by {*q_j_, μ_j_, σ_j_*}, we have

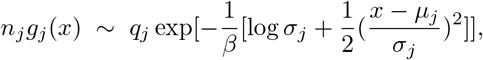

so that the tuning curve *g_j_*(*x*) must itself have a Gaussian shape (with standard deviation *β*^1/2^*σ_j_*) for each *j*. If we define the tuning curve width as the length of the interval of values 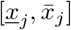 over which 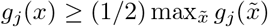, then the tuning curve width for population *j* will equal 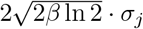.

## 4 Additional Figures for Section 3

In this section we provide additional figures to complement the main analysis in section 3. Each figure is computed using the same stimulus distribution 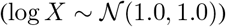 and memory size as in section 3. First, Figure 7 displays the relationship between the discrimination thresholds and the degree of stochasticity. Figure 7 validates the claim that the sudden increases in the discrimination thresholds align with transitions between adjacent categories, especially for lower values of *β*.

**Figure 7:**
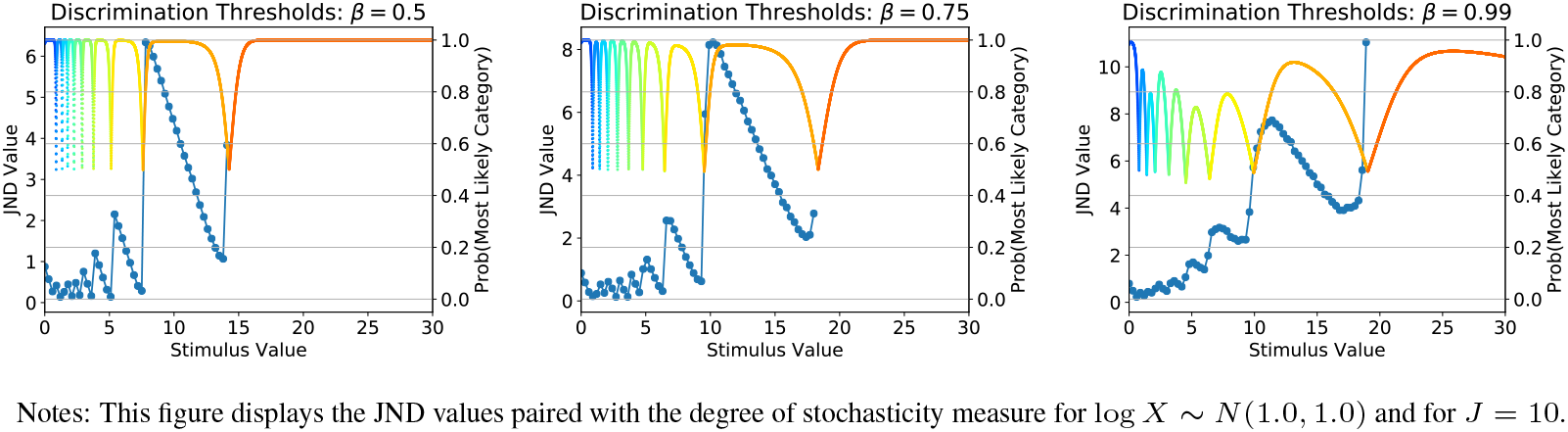
Discrimination Thresholds and Degree of Stochasticity, *J* = 10

Figure 8 displays the resulting marginal approximation and corresponding encoder and decoder for the log normal distribution considered in the main text and with *J* = 10, but considers alternative *β* values to those considered in the main text. Figure 9 and Figure 10 display the marginal approximation and corresponding encoder and decoder for the same *β* values considered in the main text {0.5,0.75,0.99} but for *J* = 15 and *J* = 20 respectively. Overall, these figures further validate the qualitative patterns described in the main text.

**Figure 8:**
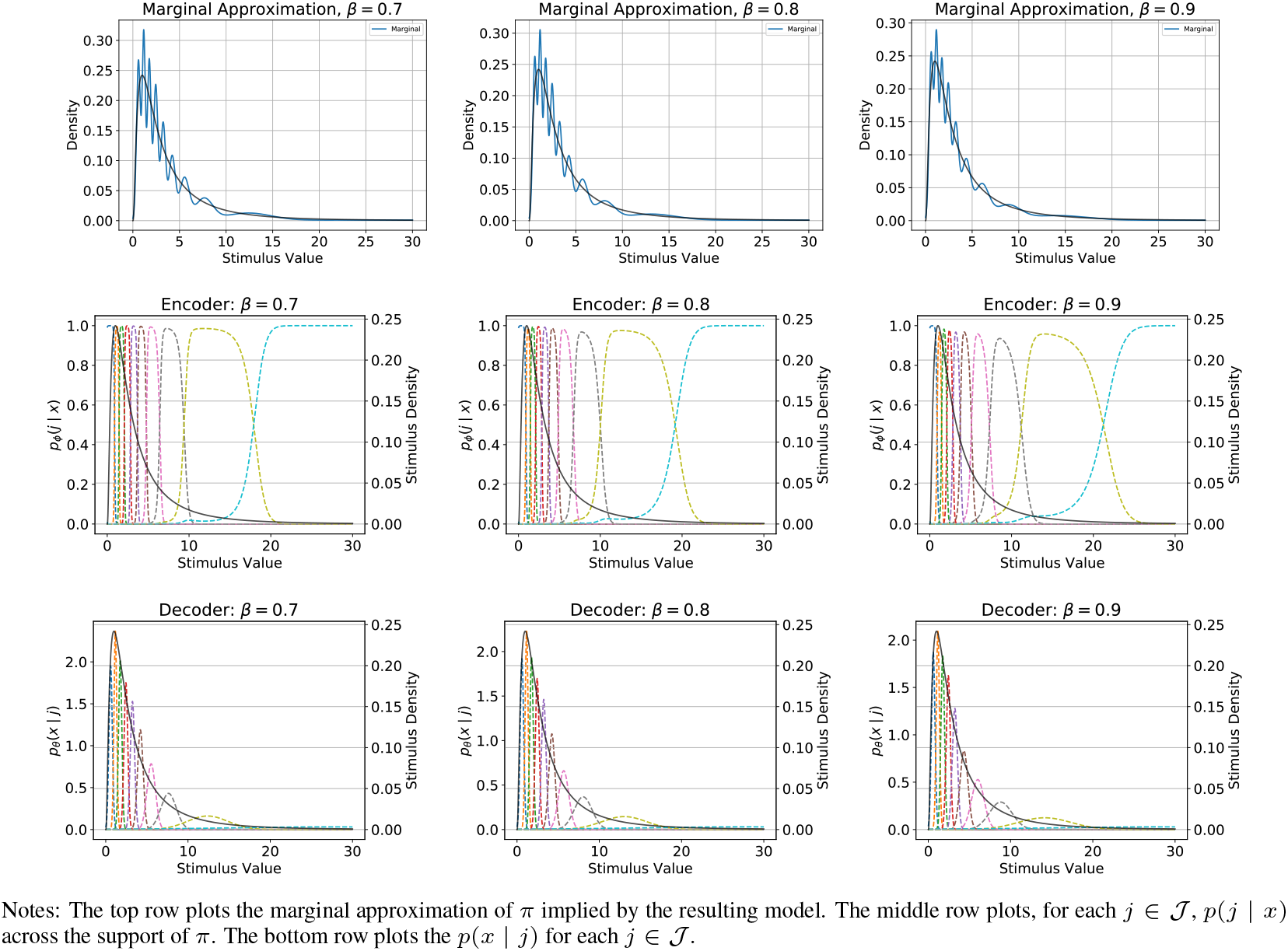
Marginal Approximation and Encoder / Decoder for *J* = 10, varying *β*

**Figure 9:**
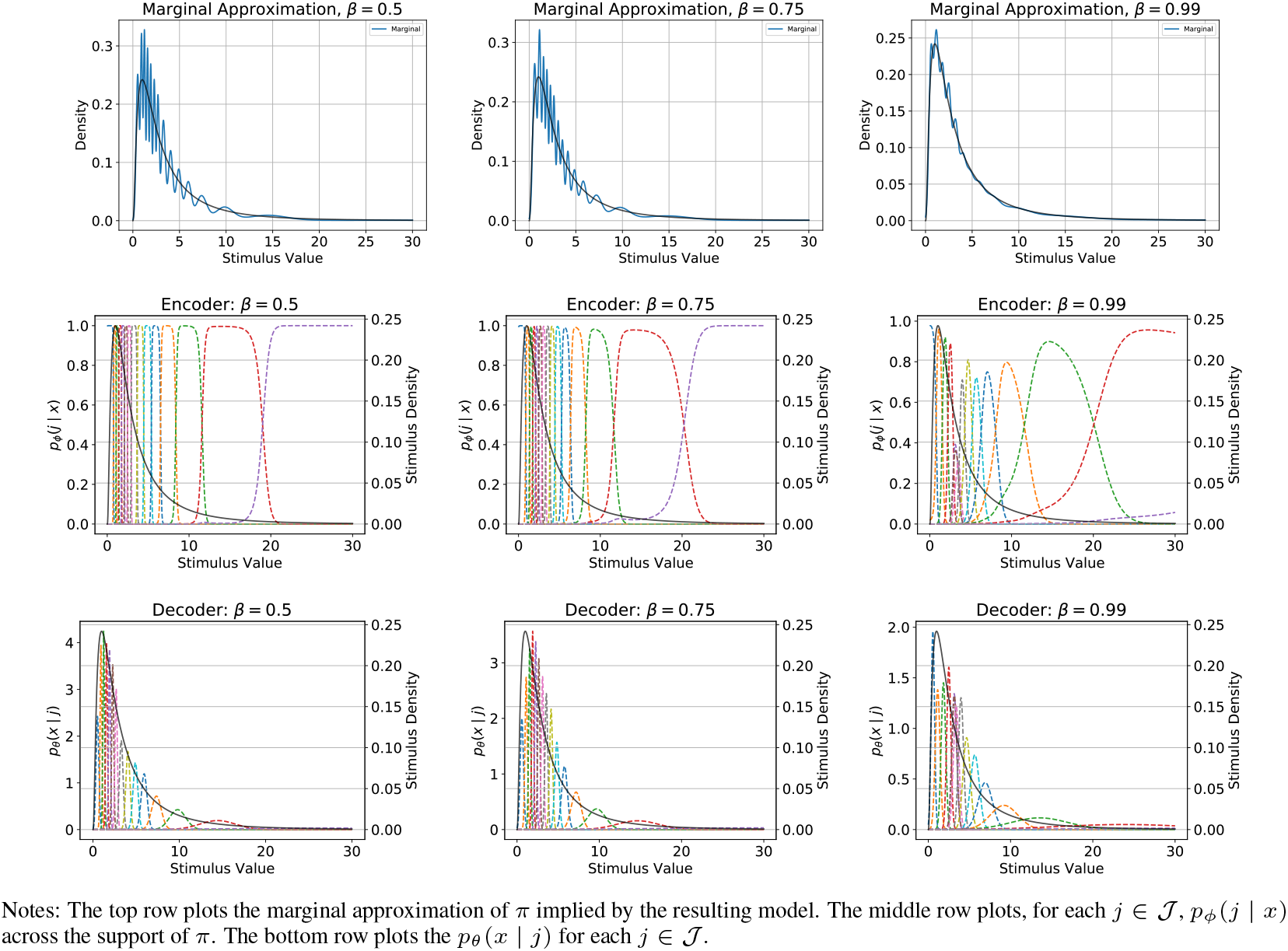
Marginal Approximation and Encoder / Decoder for *J* = 15, varying *β*

**Figure 10:**
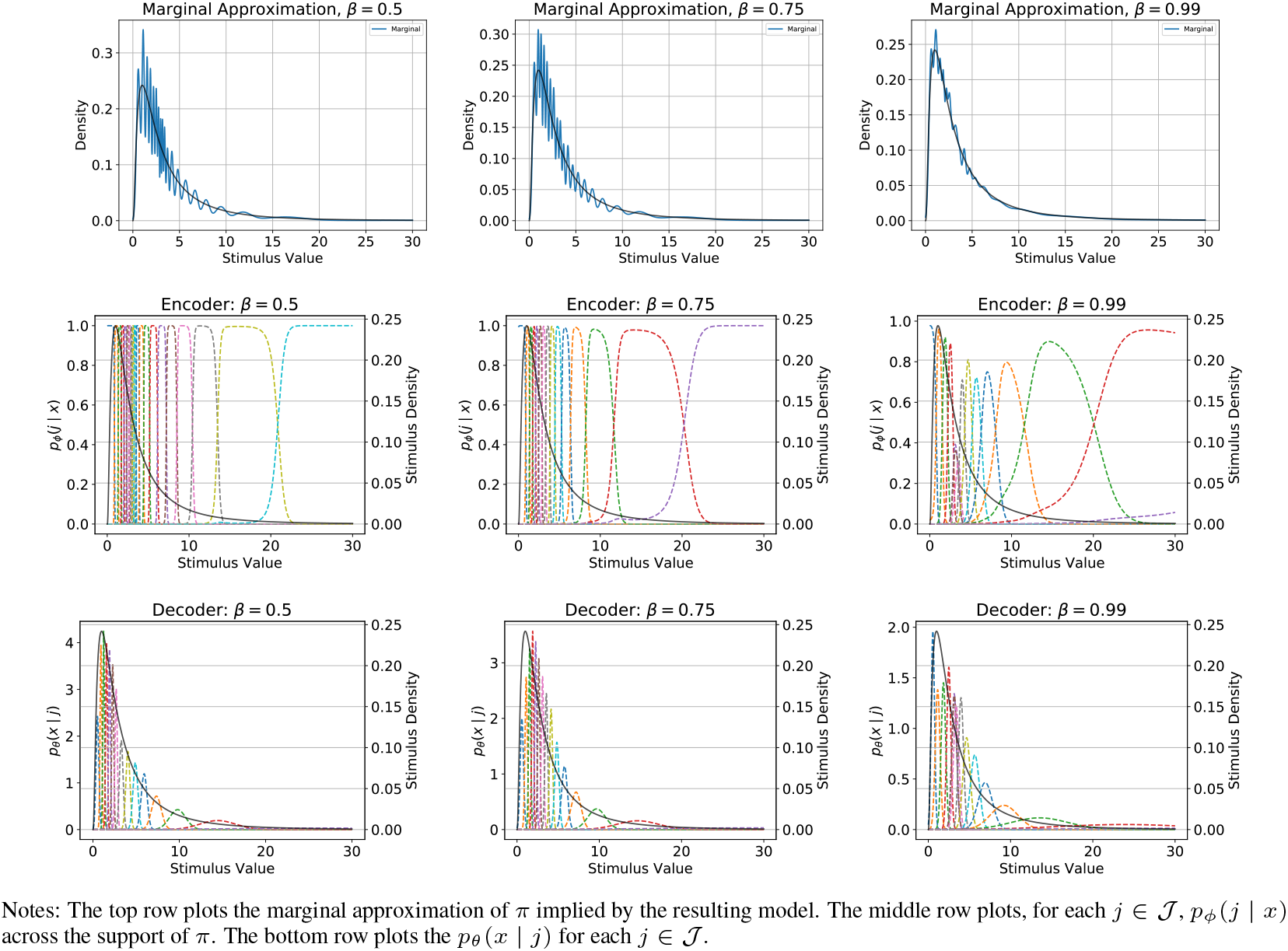
Marginal Approximation and Encoder / Decoder for *J* = 20, varying *β*

**Figure 11:**
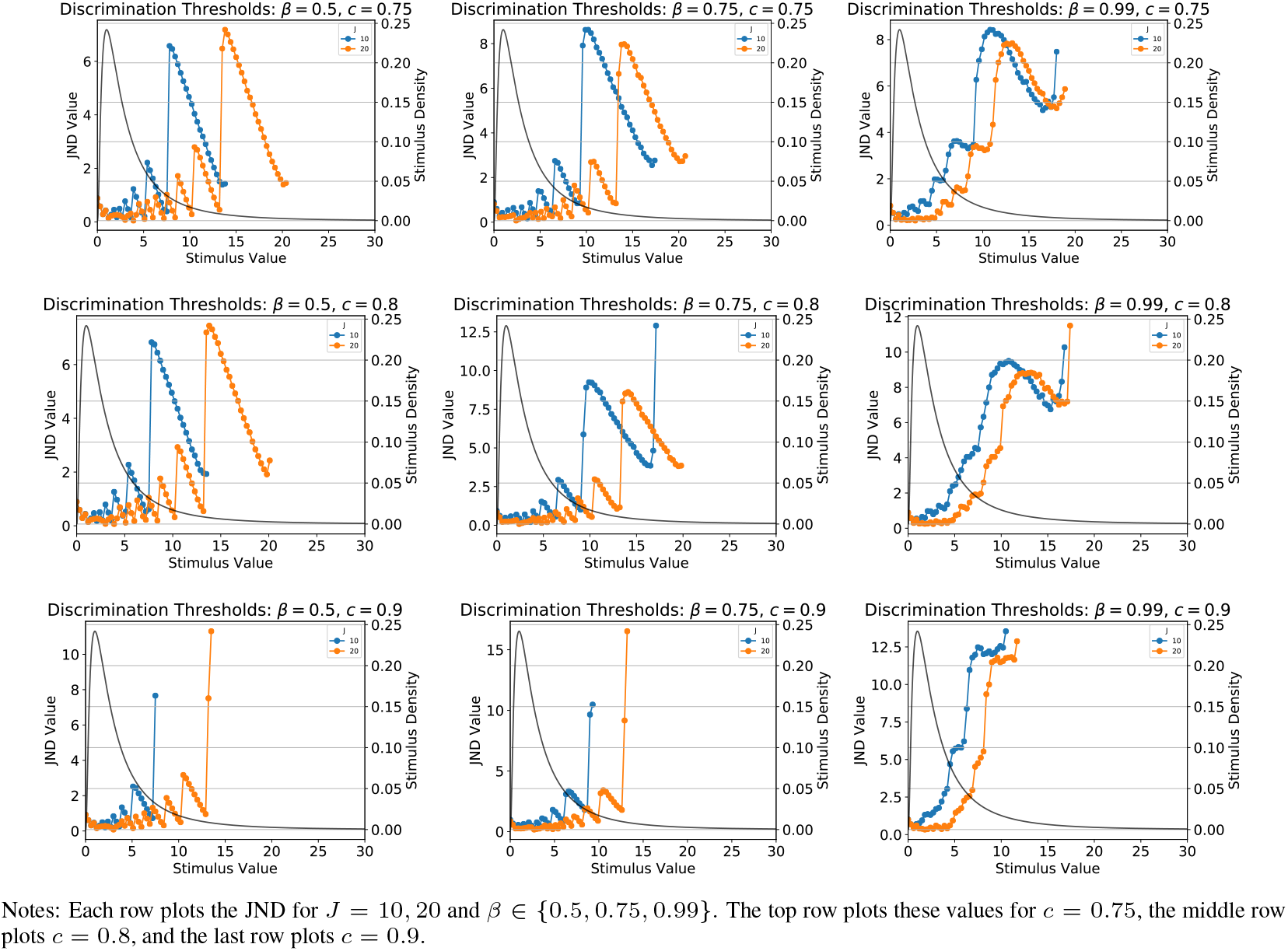
JND Plots for *c* = 0.75,0.8,0.9 and *J* = 10,20, varying *β*

## 5 Additional Figures for Section 4

In this section we provide additional figures for the calibration section. Figure 12 displays the resulting encoder, decoder, and implied marginal distribution for the calibrated parameters (*β* = 0.62, *J* = 25).

**Figure 12:**
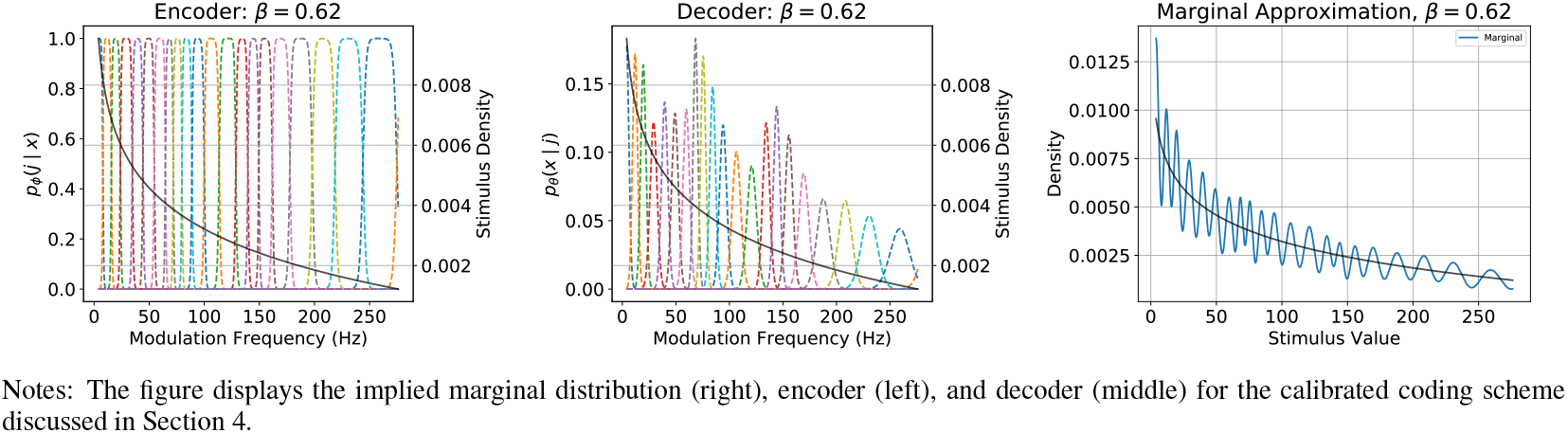
Encoding and Decoding Distribution Plots for Calibrated Discrimination Thresholds

## 6 Additional Figures for Section 5

In this section we provide additional figures for the adaptation section. Figure 13 shows adaptation from 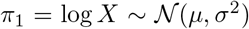 to 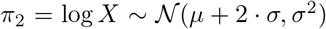 for *β* = 0.75. Figure 15 shows adaptation from 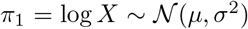 to 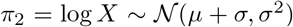 for *β* ∈ {0.5,0.75} and further illustrates rapid adaptation to *π*_2_. Finally, Figure 14 shows adaptation from 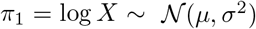 to 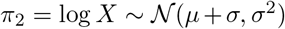 for a significantly smaller memory size. Adaptation occurs more rapidly, but the resulting parameter estimates are considerably more noisy.

**Figure 13:**
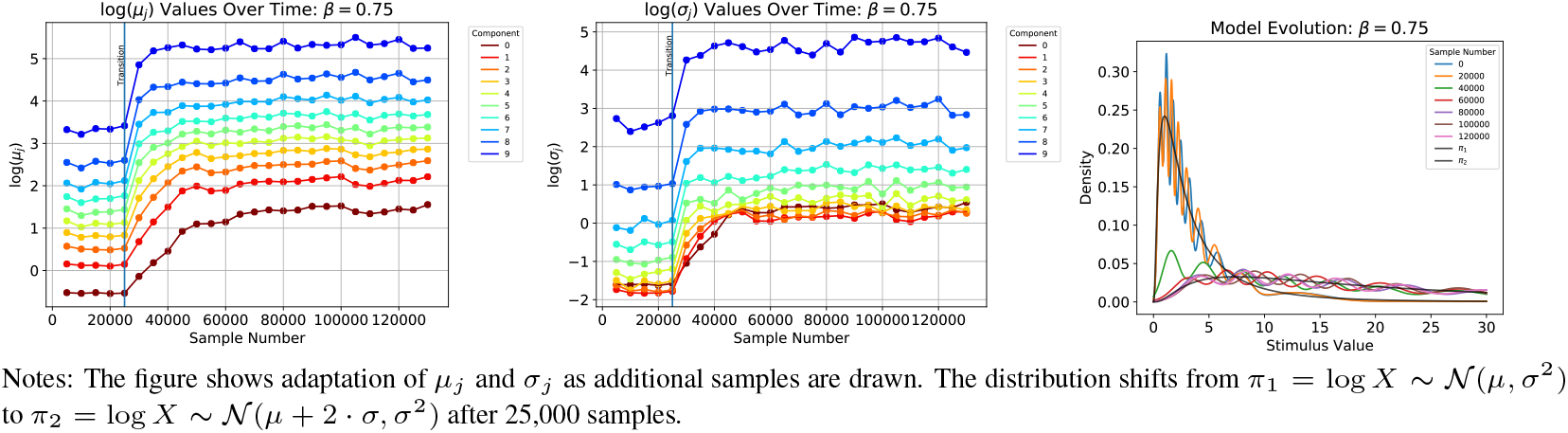
Adaptation of Parameters and Model, 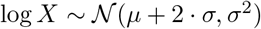, *m* = 10,000

**Figure 14:**
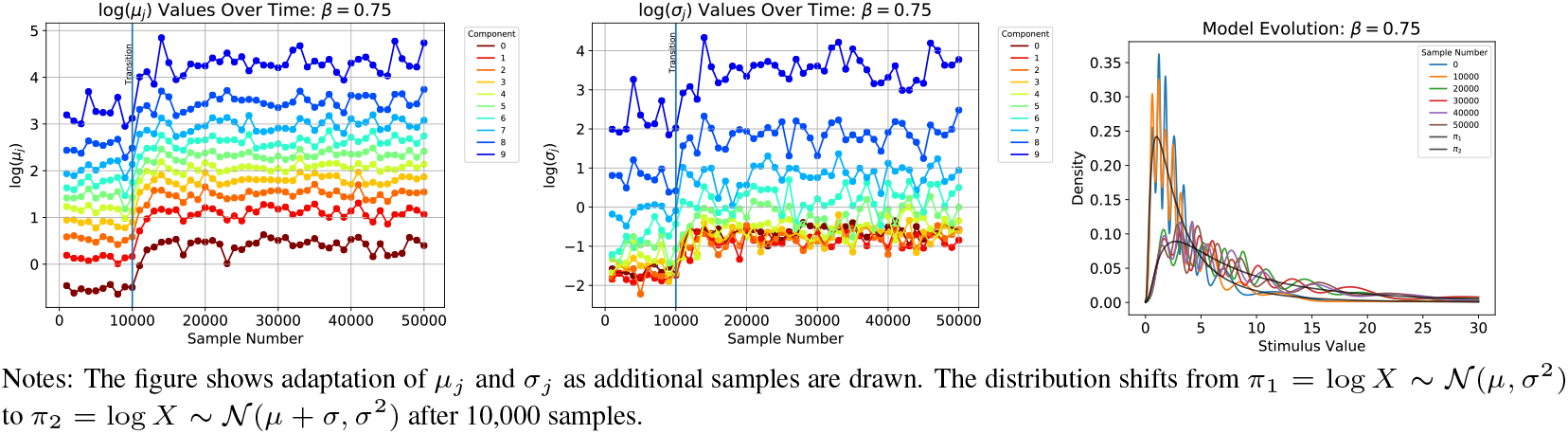
Adaptation of Parameters and Model, 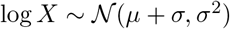, *m* = 1,000

**Figure 15:**
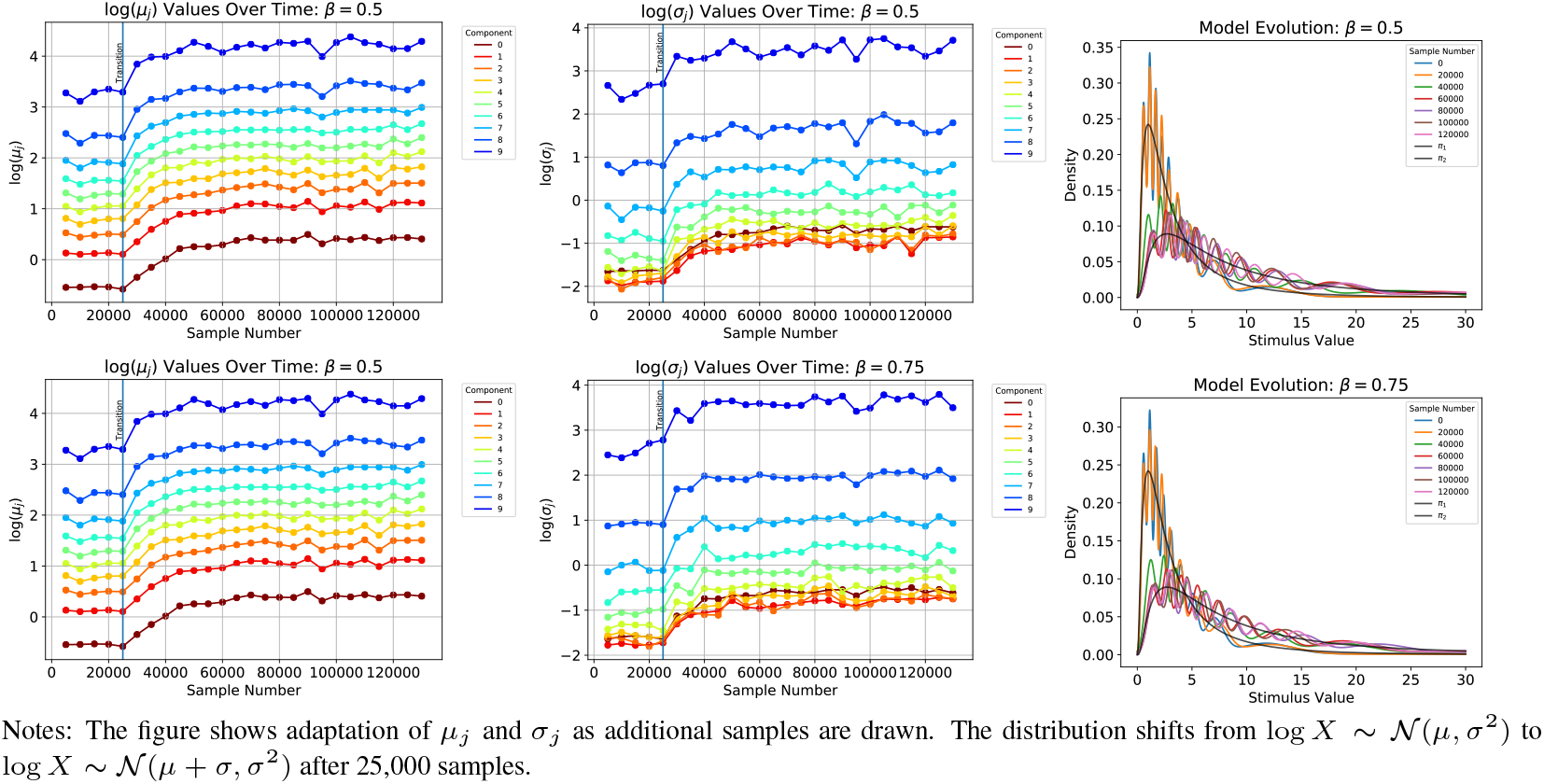
Adaptation of Parameters and Model, 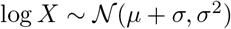, *m* = 10,000

## Footnotes

1 The details of the algorithm are displayed as Algorithm 1 in the supplementary material and the proof of its correctness is in [1]. Section 2 of the supplementary material further provides exercises demonstrating the nature of adaptation under this sampling scheme.

2 As noted by [13], poor initializations in the usage of the E/M may lead to convergence to “bad” local maxima. In order to ensure convergence, we first run the E/M algorithm initializing the *μ_j_* values utilizing k-means. Then, we re-run the E/M algorithm with random initializations until the value of the objective function converges. Our numerical experiments showed that convergence occurs after 200 random initializations for the reported values of *J*, so that we utilize this for the results that follow.

3 Section 1 of the supplementary material reports similar exercises for different stimulus distributions. In addition, section 4 of the supplementary material considers alternative values of *β* and *J* than those considered here and shows the robustness of the qualitative patterns we document.

4 We report *c* = 0.71 in the main text since this is the value of *c* for which we report the calibration results in section 4. The supplementary material provides qualitatively similar plots for alternative values of *c*.

5 Section 4, Figure 7 in the supplementary material provides evidence for this hypothesis.

6 The neural tuning width data come from [22] and the data for perceptual discrimination thresholds come from [10,29].

7 The three loss functions that we consider are: *ℓ*(*x, y*) = |*x* − *y*|, *ℓ*(*x, y*) = (*x* − *y*)^2^, or 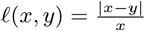. These alternatives allow for comparisons based either on the absolute magnitude of the errors or the percentage deviation; the latter case allows a loss function that is independent of the magnitude of empirical discrimination thresholds.

8 In the supplementary material we also include an exercise showing how the memory size impacts the speed of convergence of the empirical distributions.

